# Less is more: selection from a small set of options improves BCI velocity control

**DOI:** 10.1101/2024.06.03.596241

**Authors:** Pedro Alcolea, Xuan Ma, Kevin Bodkin, Lee E. Miller, Zachary C. Danziger

## Abstract

We designed the discrete direction selection (DDS) decoder for intracortical brain computer interface (iBCI) cursor control and showed that it outperformed currently used decoders in a human-operated real-time iBCI simulator and in monkey iBCI use. Unlike virtually all existing decoders that map between neural activity and continuous velocity commands, DDS uses neural activity to select among a small menu of preset cursor velocities. We compared closed-loop cursor control across four visits by each of 48 naïve, able-bodied human subjects using either DDS or one of three common continuous velocity decoders: direct regression with assist (an affine map from neural activity to cursor velocity), ReFIT, and the velocity Kalman Filter. DDS outperformed all three by a substantial margin. Subsequently, a monkey using an iBCI also had substantially better performance with DDS than with the Wiener filter decoder (direct regression decoder that includes time history). Discretizing the decoded velocity with DDS effectively traded high resolution velocity commands for less tortuous and lower noise trajectories, highlighting the potential benefits of simplifying online iBCI control.

---

An intracortical brain-computer interface (iBCI) uses spiking information from simultaneously recorded neurons to infer, or “decode”, a paralyzed user’s motor intent so it can be realized by an assistive device physically^1–4^ or digitally^5–10^. Decoders are typically built by regressing imagined or observed action against the simultaneous activity of recorded neurons^11–13^. Virtually all cursor-control iBCIs use *continuous velocity* decoders, mapping the neural activity into a 2D velocity command vector. We hypothesized that constraining decoded velocity commands to a small set of selections would improve iBCI control because the simpler repertoire of resulting cursor kinematics would be easier to learn, the discrete nature of the commands would act as a de-noising step, and stopping would be made simpler by including a zero-velocity selection. To test this hypothesis, we created the discrete direction selection (DDS) decoder, which uses neural activity to select from a small menu of cursor velocity commands. Using the jaBCI (a neural firing rate synthesizer allowing noninvasive, real-time, human-in-the-loop use of iBCI decoders^14^), we compared closed-loop performance of human subjects using DDS to three commonly used continuous velocity decoders: the velocity Kalman filter (vKF^15^), ReFIT^16^, and the so called “direct regression decoder with assist”^17^ (DR-A, an affine map from neural activity directly to continuous velocity commands). We also compared iBCI cursor control performance by a monkey using DDS and a Wiener filter (another direct regression-based decoder).

The vKF^5,11,15,18^ estimates intended velocity through a compromise between a) what it has seen the cursor do in the past, given the current cursor state (state transition model) and b) how similar the recorded neural activity is to what was expected given the predicted cursor state (measurement model). Both the state and measurement models are calibrated by regression on reference activity, and together they output a command from the continuous velocity plane. The ReFIT decoder^7,16^ uses the vKF architecture but extends the calibration procedure. The two most salient changes are that a) the state transition model predicts that the cursor obeys physical laws of movement with damping, and b) the measurement model is calibrated using the cursor states we assume the user intends rather than the actual executed states, which include targeting errors and jitter. These enhancements were made in part to overcome the difficulty vKF users had stopping the cursor. DR-A linearly maps neural activity directly into continuous velocity commands^17^. This circumvents the need for inverse solutions predicting neural activity from cursor state (that arguably introduce speed biases and errors when generating stop commands^17^), which are required by the measurement models of vKF and ReFIT. A further “assisted calibration” feature was introduced that averages the preprogrammed “training” cursor trajectory with the decoded trajectory to help regularize the calibration.

The discrete direction selection (DDS) decoder maps neural activity into a selection from a small menu of possible cursor velocity commands. Using the same training data as do the continuous velocity decoders, DDS fits a multinomial model that maps the neural activity into the probability the user is choosing each of the possible selections. We implemented DDS with nine possible selections: up, down, left, right (each with a slow and fast speed), and stop. We also included a post-prediction step that blended selections together based on their estimated probabilities, creating a smooth transition between selections. The structure of DDS was designed, in part, to simplify what is in essence a motor learning problem faced by iBCI users: the mismatch between decoded and actual intentions makes using an iBCI more like learning a new skill than it is having one’s mind read^12,19–24^. Discretizing the command space reduces high-frequency jitter in the cursor trajectory because only neural activity changes that cross the selection threshold affect cursor velocity, which may make learning iBCI control easier. Importantly, because DDS is discrete in velocity, as opposed to directly selecting target positions^25,26^, subjects can still navigate the entire workspace. The question we addressed in this study was whether trading fine-grained continuous control for the simpler and less variable trajectories DDS produces improves online performance.

We compared center-out cursor control performance in 48 able bodied naïve human subjects, each of whom used either the vKF, ReFIT, DR-A, or DDS decoders across four visits using the non-invasive “jaBCI model” which we developed earlier^14^. The jaBCI converts a person’s hand kinematics into synthetic neural activity that can be decoded in real-time with any iBCI decoder. This creates a non-invasive method to test large numbers of human subjects in closed-loop. DDS users had the top performance by a substantial margin. ReFIT outperformed vKF and improved stopping, consistent with a report in two monkeys^16^, and DR-A outperformed ReFIT. Motivated by the success of DDS in the jaBCI, we had a monkey use an iBCI with both the DDS and Wiener filter^27–29^ decoders (a common dynamic linear filter). In those experiments, DDS substantially outperformed the Weiner filter during interleaved 3.5 min blocks across seven recording sessions. The jaBCI model and monkey iBCI results together indicate that DDS can improve online performance compared to several of the more common continuous velocity decoder designs for iBCI.

## Results

### Decoder Comparison in Center-Out Cursor Control Using the jaBCI Model

The jaBCI uses a subject’s finger joint movements as input to an artificial neural network (previously trained on monkey M1 neural activity and human finger kinematics) to produce emulated neural activity in real-time (Fig.1A). Any decoder can then translate that emulated activity into commands. Each subject here was assigned one decoder. To model the daily recalibration typically required in iBCIs due to changing neurons^5,15,33–35^, at the start of each visit, subjects performed calibration posture transitions while a displayed “training” cursor^18^ followed a minimum jerk path to four cardinal direction targets. Subjects then performed an 8-target center-out task with small targets and a 0.5s hold requirement designed to be relatively difficult to avoid performance ceiling effects (see discussion for comparisons to other iBCI studies). Representative cursor trajectories are shown from a subject in each decoder group on their last visit (Fig.1B). DDS tended to generate more “staircase-like” trajectories compared to the continuous velocity decoders.

**Figure 1:**
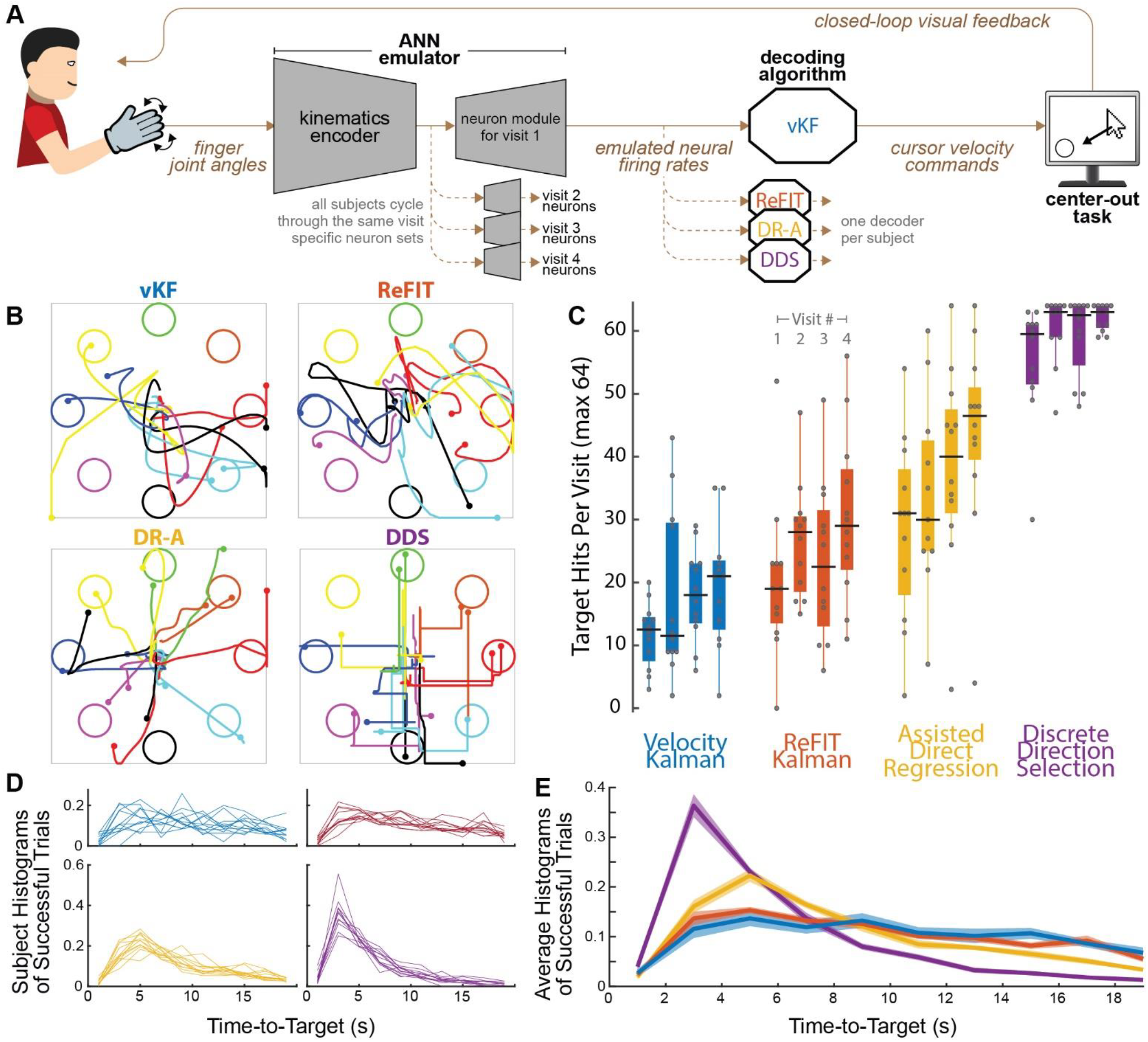
Cursor control performance of 48 naïve human subjects, each assigned one decoder for a series of four visits. A) Schematic of the joint angle BCI (jaBCI) model of iBCI use. Human subjects’ finger joint angles were input to an artificial neural network (ANN) which output emulated neural activity. Emulated neurons were changed each time a subject returned for a new visit, and the decoders were recalibrated. Subjects performed a center-out target acquisition task in real time using decoding time bins of 50 ms. B) Representative trajectories from successful trials in visit 4 from one subject in each group. Trajectories were truncated at 2.25, 2.25, 3.5, and 5.0 seconds after trial start from a visit with 17, 26, 42, and 59 total target hits for vKF, ReFIT, DR-A, and DDS decoder groups (i.e., typical success rates for each group). More time was included in DR-A and DDS to capture fuller trajectories because subjects moved more slowly overall (see also Fig. 2B). C) Boxplots of number of targets hit in the center-out task out of 64 possible, outer grouping by decoder used (color) and inner grouping by visit number. Performance increased with repeated visits and also varied with the decoder. Dots are individual subjects, black bars are the median, solid box is the interquartile range, and whiskers extend to 1.5 s.d. ± of the mean. D) Trial time histograms for each subject (successful trials only, including the 0.5 s hold time, 2 second bins). Time distributions were consistent across subjects. Intra-group variability was larger in groups where fewer targets were hit, in part because fewer samples were available. E) Across-subject averages (shaded patches are ± s.d.) of the histograms in D show that groups with better center-out cursor control performance (C) hit targets more quickly (more distribution mass at lower times).

The average number of targets hit differed between decoder groups, 16.7, 25.2, 36.1, and 59.4 for vKF, ReFIT, DR-A, and DDS (Fig.1C ANOVA F(3,175)=131, p<0.001, all post-hoc comparisons p<0.002). There was no performance difference between the cardinal and intermediate direction targets (Fig.S1, paired t-test pooled across visits p=0.41, pooled by decoder only DR-A was significant with 1.3 more intermediate targets hit p=0.009), which is important for two reasons. First, the decoder training data were collected based on trajectories only to the cardinal targets, showing subjects could move to targets not explicitly in calibration. Second, the DDS decoder only had cardinal direction velocities available, implying that a limited set of selections does not prevent subjects from traversing the full workspace. A similar pattern of decoder performances held for a more challenging virtual keyboard typing task subjects attempted for the last 10 min of each visit (Fig.S2).

Subjects hit more targets on each subsequent visit despite the changing set of emulated neurons (slopes of linear fits across visits were 2.4-p=0.08, 2.7-p=0.06, 5.0-p=0.12, 1.9-p=0.11 hits per visit for vKF, ReFIT, DR-A, and DDS). The low learning rate for DDS was due to subjects reaching the performance ceiling. Although the emulated neurons changed each visit, subjects used the same four sets in the same order. This let us control for neural variability to compare directly between decoders in a way that an iBCI study cannot, as neuron selection is not under experimental control. After subtracting off the improvement due to learning across visits, there were no significant performance differences remaining between visits (Tab.S2 ANOVA F(3,172)=0.59, p=0.70). This indicates that the set of neurons themselves did not have a strong influence on performance, agreeing with findings in our prior work^14^.

Time-to-target was consistent among subjects within each decoder group (Fig.1D). Although DDS had a maximum speed, unlike the continuous velocity decoders, DDS subjects reached targets faster (Fig.1E). To measure stopping difficulty we used “dial-in time”, defined as the time the cursor spent within two target radii of the target center, excluding the 0.5s hold. Dial-in times were 2.1±0.6, 2.1±0.5, 1.5±0.4, and 1.1±0.3 seconds for vKF, ReFIT, DR-A, and DDS (µ±σ, all pairwise t-test comparisons were p<0.001 except vKF-ReFIT with p=0.57). Only subjects using the ReFIT and DDS decoders came to complete stops in the target (Fig.2A), and vKF and DR-A subjects would slow down or circle within the target to satisfy the hold time requirement. DR-A subjects had the lowest average cursor speeds, and all the continuous velocity decoders had a long tail of trials with very high speeds where the cursor became difficult to control (Fig.2B).

**Figure 2:**
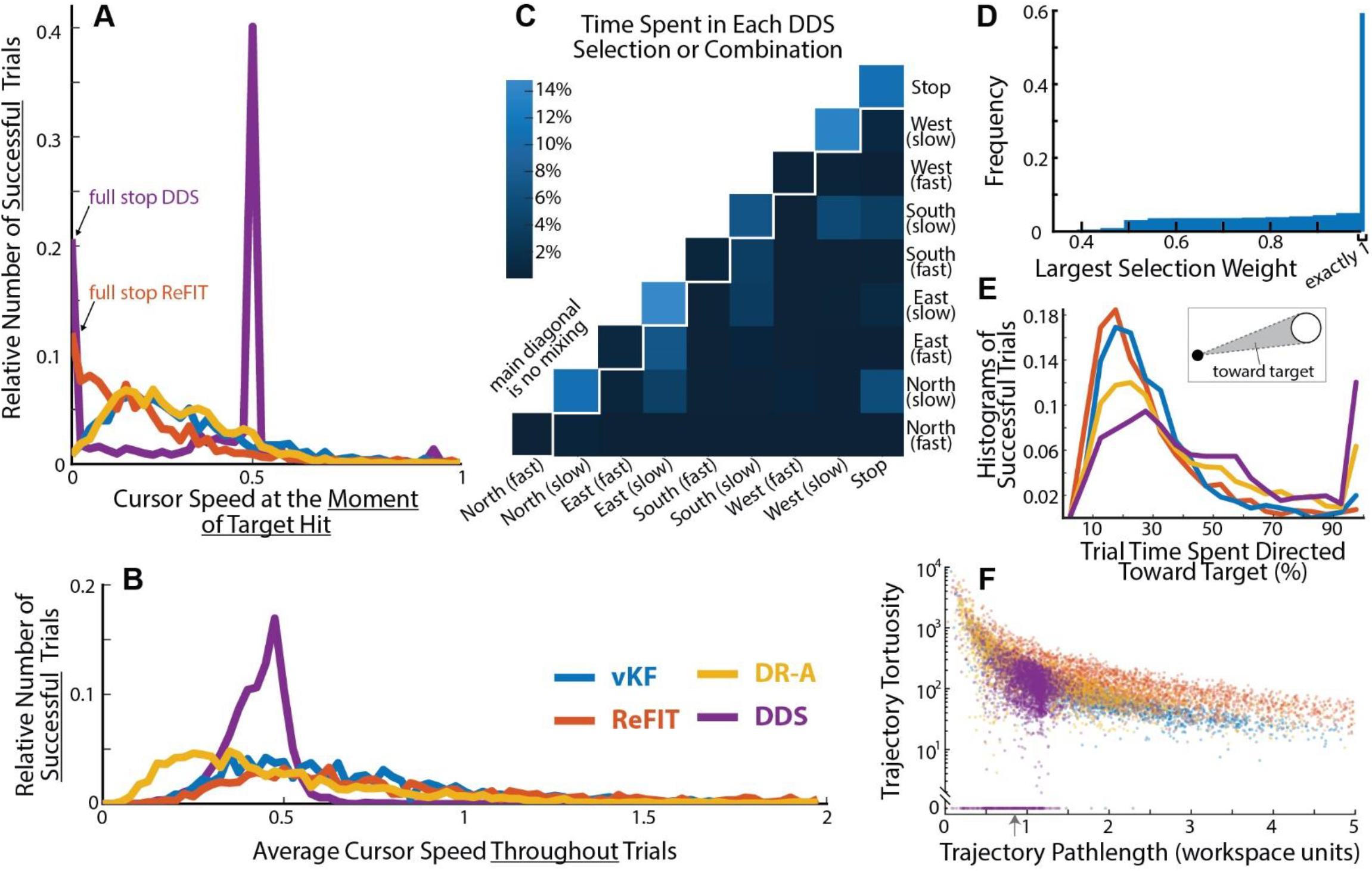
Synthesis of cursor control strategies across decoders. A) The distribution of speeds at the moment of target hit. ReFIT and DDS users often brought the cursor to a full stop, while vKF and DR-A users did not. The DDS speed selections were 0.0, 0.5, and 1.0 units/s, thus giving rise to the peaked distributions. B) The distribution of average cursor speed throughout the entire trial shows DR-A and DDS groups have fewer high-speed trials. Histograms contain 804, 1208, 1710, and 2850 trials for vKF, ReFIT, DR-A, and DDS for panels A, B, and E. Speed was measured in workspace units per second, and the workspace was a square of side length two. C) Time DDS subjects spent in each selection or combination. The four “slow”-only directions and “stop” had the highest frequencies in the data. D) The distribution of the largest selection weight shows subjects spent most of the time in a single selection (exactly 1). This panel shows the degree of mixing, whereas panel C shows whether *any* mixing occurred. E) Histograms showing the percent of each trial the cursor spent heading toward the target (i.e., a velocity within the angle subtended by lines tangent to the target originating from the cursor, inset). F) Trajectory pathlength versus trajectory tortuosity (integrated instantaneous curvature change normalized by pathlength) shows DDS subjects executed shorter and spatially simpler trajectories than other decoder groups. The occasional sharp turns in DDS trajectories generated tortuosity similar to the continuously smooth and simple arcs of vKF trajectories, although they were typically shorter. ReFIT trajectories contained the most erratic changes in direction; see also panel E and Figure 1B. The gray arrow denotes the straight-line distance from the workspace center to the center of the peripheral targets for reference.

Figure 2C shows a histogram of the percent of time DDS subjects spent in each of the 9 velocity command selections. DDS mixes selections together when their estimated probabilities are close, which is displayed in the histogram as the intersection cell of each combination of selections, with the self-intersection along top diagonal representing the unmixed selections. DDS subjects spent approximately 60% of all trial time in unmixed selections. When selection mixing did occur, it was mainly between the “slow” and “fast” velocities in the same direction or between the “slow” and zero-velocity selections (Fig.2C), with only 16.7% of time spent in selections that were mixed between different directions. To produce the final velocity command DDS averaged all selections weighted by their estimated probabilities. To understand how strongly selections were mixed (not just if they were mixed at all, as in Fig.2C), figure 2D shows the histogram of the largest selection weight at each moment. Because the weights across all selections sum to one, the greater the largest weight, the less mixing occurred. The far right of the histogram is when the largest weight was exactly one (i.e., fully unmixed selections, equivalent to summing the top diagonal cells in Fig.2C), which was by far the most common. When mixing occurred, all combinations from equal mixing (largest weight 0.5) to near complete dominance by a single selection (largest weight 0.99) were equally likely. This is shown in figure 2D by the approximately uniform distribution from 0.5 (≈3.1%) to 0.99 (≈5.0%).

DDS Cursor velocities were oriented toward the target (within the angle formed by the pair of lines tangent to the two sides of the target) more often than other decoders (Fig.2E) and DDS had substantially more trajectories that moved toward the target throughout the entire trial (sharp increase at the 100% bin). We evaluated cursor path simplicity using a tortuosity measure that integrates the instantaneous curvature derivative^27^, such that simple constant curvature shapes (an arc of a circle or straight-line segment) have zero tortuosity. We found that ≈11% of all DDS trajectories were completely straight (Fig.2F, bottom points). The remaining trajectories had lower tortuosity and shorter pathlengths than ReFIT and DR-A. Trajectories from the vKF group had tortuosity similar to DDS (subjects tended to make smoothly varying constant curvature trajectories), but with longer pathlengths. Unlike the other decoder groups, DDS subjects rarely made very short trajectories that spiraled in on themselves (upper left) or simple but long trajectories (far right) caused by high velocities.

### Decoder Comparison in Center-Out Cursor Control by a Monkey using an iBCI

To determine if the DDS performance translates from jaBCI to an iBCI, we compared the ability of a rhesus macaque to operate a brain-controlled computer cursor using a Wiener filter (WF) decoder and the DDS decoder. Following decoder calibration in each session, the monkey used the two decoders in interleaved 3.5-minute blocks, with the session’s initial decoder determined randomly (Fig.3A-B). Figure 3C shows the behavioral performance in a representative session. In most blocks, the monkey achieved a higher success rate and more rapid target acquisition with DDS than with WF. Notably, in the last four blocks, performance was maintained with DDS but dropped with WF. Figure 3D shows example cursor trajectories from successful trials. The “staircase” structure in these trajectories was also present, but less exaggerated than that of the human jaBCI subjects (Fig.1B), and correspondingly spent less time in unmixed selections (Fig.3E). Figure 3F-G compares the performance of the two decoders across all seven sessions. DDS had a higher success rate than WF in 31 of 38 pairs of temporally adjacent blocks and lower acquisition time in 26 pairs. The monkey achieved a higher overall success with DDS than WF (DDS: 61%±3% (mean±s.e.), WF: 37%±3%, p<0.001, two-tailed paired t-test). Time-to-target with DDS was slightly lower (DDS: 1.92±0.03 s (mean±s.e.), WF: 2.02±0.05s, p=0.042, two-tailed paired t-test).

**Figure 3:**
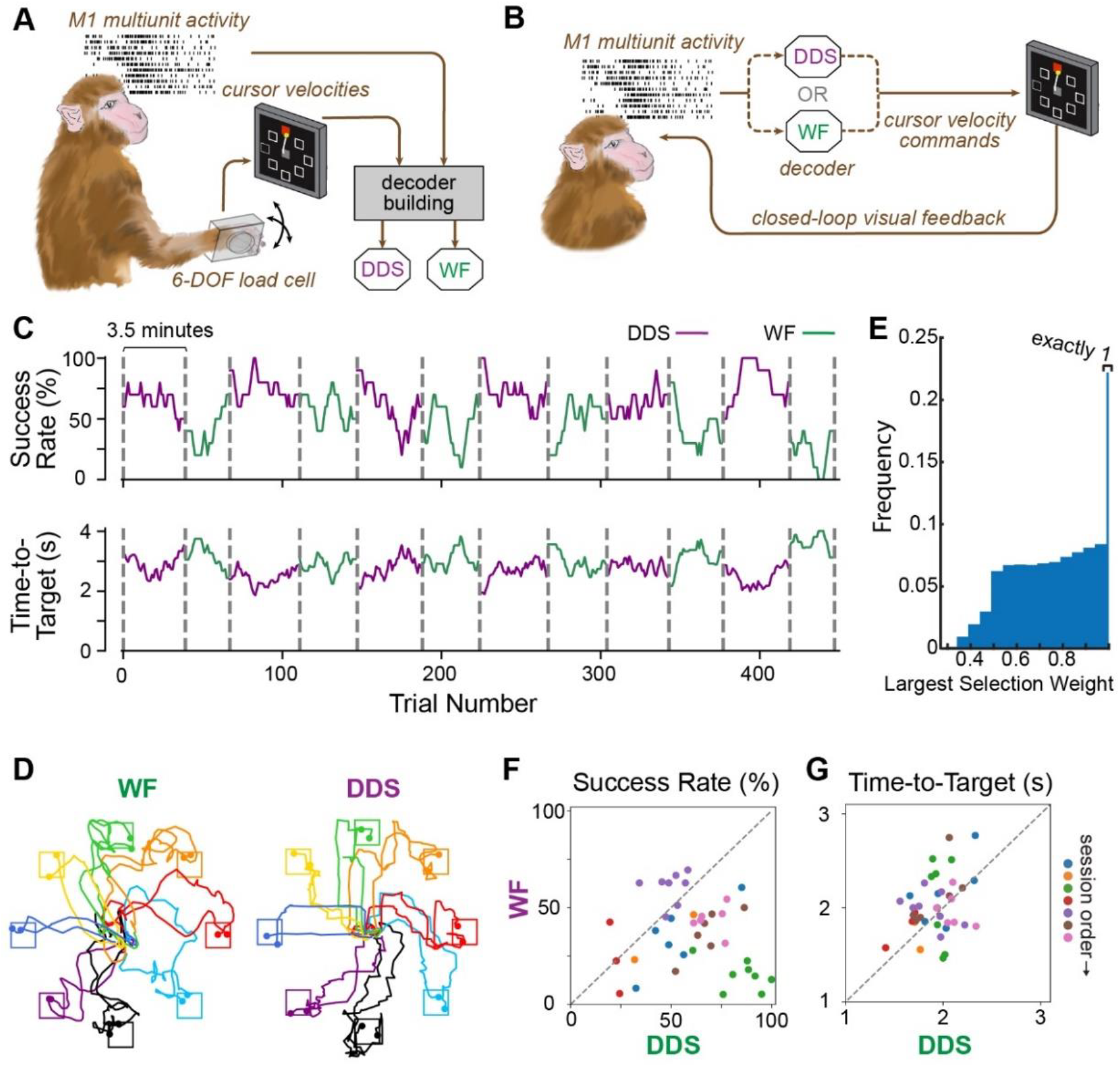
Cursor control performance of one monkey using two decoders. A) Schematic of the decoder building process. A monkey was trained to move a cursor on a monitor by exerting 2D forces on a small box placed around its right hand. M1 multiunit activity and cursor velocities were synchronously recorded and used to build Wiener filter (WF)- and DDS-based decoders. B) Schematic of the online closed-loop iBCI test. Once the decoders were built, the monkey was required to control the cursor on the monitor with M1 multiunit activity using either of the decoders. C) Behavioral performance during a representative iBCI test session, as measured by success rate and time-to-target in causal 10-trial sliding windows. The two types of decoders were tested in interleaved 3.5-minute blocks, as indicated by the vertical dashed lines. D) Representative trajectories between the go cue and the delivery of reward from successful trials, two of which are plotted for each target. E) Histogram of largest DDS selection weights over all sessions has a distribution qualitatively similar to that of the human jaBCI subjects (Fig. 2D), although spending somewhat less time in the purely unmixed selections. F-G) Comparisons of success rate and acquisition time for the two decoders across 7 sessions (indicated by symbol color). Each dot represents the comparison between two adjacent blocks from a given session.

### Offline Analysis is Not Predictive of Closed-Loop Performance in jaBCI

The jaBCI offline decoder accuracy was not predictive of closed-loop proficiency. Figure 4A shows there was no correlation between the average similarity of the calibration training cursor to the decoded trajectories and the subsequent number of hits in the center-out task (visits pooled across decoders r=-0.03, p=0.72). This indicates that a decoder’s ability to reconstruct cursor kinematics of offline training data is not a key determinant of its success online.

**Figure 4:**
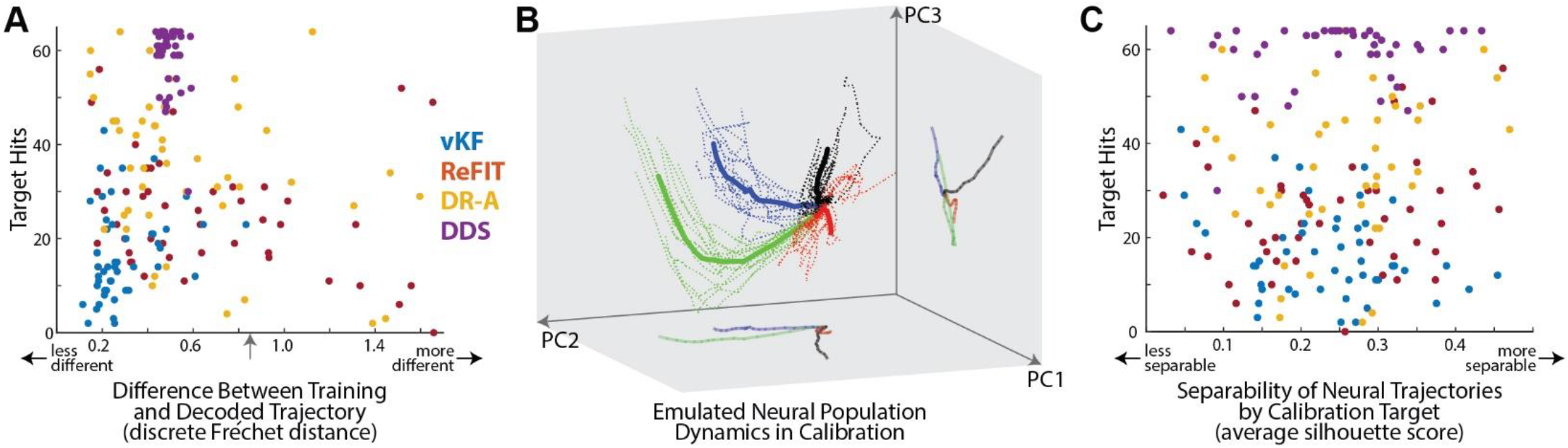
Accurate offline decoding in jaBCI was not predictive of closed-loop performance in the center-out task, nor was the low-dimensional separability of the emulated neural population dynamics. A) There was no correlation between the difference of decoded and training trajectories (abscissa) and online control proficiency (ordinate). Trajectory similarity was measured with the discrete Fréchet distance, where zero indicates identical trajectories and grows with increasing dissimilarity. The gray arrow denotes the trajectory difference that would result if the decoded cursor had stayed at the origin throughout the trial for reference. B) A typical example of emulated neural population dynamics during calibration from one subject visit. The low-dimensional space is defined by the top three PCs of all emulated neural activity during the calibration. Individual trials are dotted curves, colored by the calibration target for that trial, and solid traces are the spatial average over all trials to that target. “Neural trajectories” typically formed distinct paths through PC space according to target and originate from the same location (waiting for the trial go cue). C) We measured separability in terms of the silhouette score, which can range from −1 to 1, where larger values indicate that trajectories to a given target are increasingly similar to each other relative to trajectories to other targets. The separability of emulated neural population dynamics (average silhouette score for all trials in a subject visit, abscissa) was not correlated with online control proficiency (ordinate).

The evolution of neural activity through neural state space during calibration trials was typically distinct for different targets (Fig.4B, projection onto the top three PCs capturing 97%±3% variance). However, the degree of separability was also not predictive of closed-loop proficiency. Figure 4C shows no correlation between the separability of projected neural population trajectories and the number of targets hit (visits pooled across decoders r=0.08, p=0.26). The lack of a relationship between neural trajectory separability and performance is surprising from a purely machine learning perspective, where accurate classification depends on being able to partition data in the feature space^28^, as well as from a neuroscience perspective where some hypothesize that separability of low-dimensional neural population trajectories fosters more robust computation^29–33^.

## Discussion

The discrete direction selection (DDS) decoder maps neural activity into a selection from a short list of velocity commands, and outperformed traditional continuous velocity decoders in the human-in-the-loop jaBCI model (N=48) and in iBCI control by a monkey. The result opens the door to a new type of decoder that exploits the accuracy of discrete selection^25^ without sacrificing the flexibility of full workspace accessibility. The work also highlights the utility of the jaBCI, which can generate predictions about decoder performance that are statistically robust across subjects to screen ideas for follow-up in the more invasive and resource-intensive human or monkey iBCI studies.

Simultaneously decoding continuous cursor direction and speed is challenging. The stochasticity of neural activity leads to noise in decoded cursor velocity (and thus position), which degrades the ability to navigate to and hold the cursor still inside a target. Information about intended speed may also not be well represented in the cortical populations typically used in iBCI^34^, making it harder still to decode intended stopping. ReFIT addressed this by setting velocity to zero (regardless of decoded velocity) whenever the cursor is inside a target during calibration^16^, which allowed jaBCI subjects to stop consistently inside targets (Fig.2A). The speed dampening KF decoder (not tested here) dynamically scales down the speed prediction in the KF state transition model when changing directions^34^. Some argue that affine transforms directly from neural activity to velocity commands, like DR-A and WF, improve cursor speed estimates over KF implementations that estimate neural activity from cursor states^17^. However, DR-A subjects did not stop the cursor (Fig.2A), but DR-A did prevent many trials with high, and perhaps unmanageable, average cursor speeds (Fig.2B).

Another approach has been to use two parallel decoders, one responsible for continuous velocity control and the other for a “neural click” command. Decoding a discrete state propagates far less of the neural variability into cursor state noise because the binary decision will not often flip in response to small changes in the neural input. Clicks have been implemented with Bayes^8^, LDA^35^, or hidden Markov models^36^ or by blending a zero-velocity command with the continuous velocity command in proportion to an LDA classifier’s confidence in the click state^37^. The continuous velocity decoder can even be abandoned altogether for accurate and efficient target selection from preset grids^25^ or letters^26^; however, this approach forecloses the possibility of continuous control.

The premise behind DDS is to use the noise-rejection advantage of discrete classification to directly select cursor velocity while retaining the ability to traverse the entire workspace, since position is unconstrained. Unlike methods that use smoothing to denoise trajectories (e.g., long bin widths or temporal averages^38,39^), DDS does not introduce lag because the velocity commands are issued every time bin. The noise rejection comes at the expense of coarsely decoded velocity (e.g., staircase-like trajectories, Fig.1B and 3D), but this tradeoff appears to benefit performance (Fig.1C and 3F) and create simpler trajectories (Fig.2F). Correspondingly, the coarse-grained commands led to below average offline accuracy (Fig.4A) because discrete velocities could not reconstruct the continuously varying calibration trajectories (consistent with the observation that offline accuracy is not predictive of closed-loop iBCI performance^13,39–45^). The extra time incurred when taking a staircase rather than straight trajectory to the target is small, so accepting coarse decoding for noise rejection appears to be a good tradeoff in tasks where trial times longer than a second are permissible. For example, in our cursor task a staircase is 176ms longer than a straight path to the target (assuming the slow velocity selection), <2% of the average trial time. In fact, subjects tended to spend more time in single, unmixed selections as their performance improved across visits (54%, 57%, 63%, and 67%), meaning they did not appear to avoid the staircase trajectories.

The selection mixing in DDS softens the boundaries between discrete velocity commands, which makes those regions more similar to continuous velocity control and thus permits more cursor noise. The amount of mixing can be tuned from winner-take-all, *γ* = 1/2, to an average weighted exactly by the decoded probability estimates, *γ* = 1. We used *γ* = 0.85, but future studies could optimize *γ* to improve performance. Anecdotally, we found that *γ* = 1/2 was very difficult to control, perhaps because without any mixing there were no visual cues for when the decoder was about to switch selections, whereas with mixing there is a short and smooth transition between velocity commands as the neural activity approaches the classifier’s selection boundary. Optimizing other DDS parameters, such as the number of selections, the speeds of each selection, access to past neural activity, the classification model, or even the calibration protocol (e.g., adding separate calibration trials for different cursor speed selections), may improve performance.

The reason for the counterintuitive performance gain that came from coarse-graining velocities is unclear. Speculatively, it may be that DDS effectively “teaches” subjects to execute commands with coarse precision (e.g., more like “go left” than “speed x cm/s at heading y degrees”). This may improve closed-loop control because decoding a selection from a small set of options can be done more accurately than can high-precision velocity commands, thereby eliminating the need to correct persistent errors in heading generated in continuous velocity decoding. In essence, DDS does not attempt to decode commands at higher precision than the neural activity supports. Future work could investigate how the supported precision (i.e., number of available selections) depends on channel noise or number of recorded neurons.

This study adds to the body of evidence of the validity and utility of the jaBCI model for decoder evaluation. We corroborated two findings shown in iBCI studies: 1) ReFIT’s ability to improve stopping^16,46^ and 2) ReFIT’s better performance than vKF^16^. To our knowledge there have been no direct iBCI comparisons between DR-A (or DR-like decoders) and ReFIT, so DR-A’s better performance in the jaBCI is a prediction rather than a validation. We also found that DDS outperformed DR-A in the jaBCI and a similar direct-regression decoder (WF) in a monkey iBCI. This is strong evidence that decoders designed, tested, and found to perform well with the jaBCI will have correspondingly strong performance in iBCIs. jaBCI subjects using continuous velocity decoders hit fewer targets than what is often reported for iBCIs, in part because we used smaller targets (1.7% of workspace area rather than 9.3%^5^, 5.7%^16^, or 3.2%^18^) that were farther from the center (35% of the total workspace distance to the nearest target edge rather than 16%^5^, 26%^16^, and 32%^18^). In an iBCI task of comparable difficulty (1.2% area and 37% distance)^6^, better performance was reported after 1000 days, as opposed to four in this study. In prior jaBCI validations we matched the iBCI task parameters precisely, and observed equivalent performance in jaBCI^14^. The jaBCI-emulated neural activity also matched firing rate statistics and low-dimensional population behavior of monkey M1 activity during reaching ^14^. Here we opted for a more difficult task to prevent subjects from reaching a performance ceiling, which would compress differences between decoders, although DDS subjects appeared to reach this performance ceiling anyway (Fig.1C).

These results demonstrate that an iBCI decoder using coarse-grained velocity commands can outperform continuous velocity decoders. Giving up the precision control of continuous decoding and accepting worse offline performance to achieve simpler low noise cursor trajectories appears to be a beneficial trade for online control. This reinforces the importance of closed-loop subject-decoder interactions in iBCI design and control.

## Methods

### Human Subjects

Studies were approved by the Florida International University IRB. Inclusion criteria were neuralogically intact, able bodied people ages 18-65, without injury affecting manual dexterity, whose hand could fit in the CyberGlove III (CyberGlove systems). Subjects were compensated $10/visit. A total of 56 subjects with no knowledge of the experiment were recruited. Eight subjects were excluded post-recruitment: one for failing to complete all visits, two for having fingers too short to reach the distal glove sensors, and three due to technical errors during data collection. The remaining 48 subjects (24 females, 24 males, aged 18-42) participated in four visits within a 14-day period. On the first visit, subjects completed a brief demographic survey and were pseudorandomly assigned to one of the four decoder groups. At the start of each visit subjects sat approximately 70 to 90 cm in front of the computer monitor (34 cm X 60 cm) and an experienter helped them don the CyberGlove.

### The jaBCI Model

The jaBCI model and its validation using the vKF decoder is described in detail in Awasthi et al. 2022^14^ and illustrated schematically in figure 1A. Briefly, a subject wears a CyberGlove III that monitors the relative excursions of 19 different hand and finger joint angles. The same pre-trained artificial neural network used in the jaBCI validation study ^14^ computes a set of emulated neural firing rates based on the preceeding 100ms of joint angle input data. To align with contemporary firing rate integration bin widths used in iBCI studies, we set the emulator to emit neural firing rates every 50 ms. At each time bin the subject’s decoder uses the emulated neural activity to compute the velocity of the computer cursor and the graphical display is updated accordingly. The joint angle tracking, neural emulation, and decoding all run in real-time. To mimic the phenomenon of neuron turnover in iBCI studies^18,47^, we used four separate neural readout modules (one for each visit) that emulated 71, 45, 70, and 82 neurons, thus requiring subjects to recalibrate their decoders at the start of each visit. Since we had previously found the effect of neuron set on performance was neglidgible, we selected the sets used by the most subjects in the validation study^14^.

### Protocol for jaBCI Use

The jaBCI emulated a different set of neurons each visit (Supplementary Fig. 3A - such that all subjects cycled through the same four sets on subsequent visits); accordingly, the subject’s decoder was calibrated at the start of each visit (Supplementary Fig. 3B). To do this, subjects made a fist posture that was comfortable for them and were then shown a picture of a hand posture in the center of the workspace and a corresponding target in one of the four cardinal directions. Subjects were instructed to initiate a transition from the fist to the displayed posture when the target changed color, following a computer cursor traveling via minimum jerk trajectory between the workspace center and the target (i.e., a “training cursor^18^”). They were instructed to complete the transition in 1.2 seconds, just as the training cursor came to rest in the target. The four targets were presented in sequence and the sequence was repeated twice, consisting of one block, then the decoder was recalibrated using the data from the current and all prior blocks. Users performed 7-9 blocks. In blocks 1 and 2 only the training cursor was displayed, while in all subsequent blocks the decoded cursor was also displayed. Individual calibration trials were flagged as outliers automatically, using PCA through time to project each trial to a point in a 6-dimensional latent space. Each 6D projected point was labeled according to the calibration target displayed during the trial, and we determined each trial’s similarity to other trials to that target by computing its silhouette score^48^. Trials were flagged as an outlier (and removed from the decoder training data) if the silhouette score was less than zero, i.e., the given trial was more similar to trials from another calibration target than to its own. The silhouette score used in the outlier detection was based on the Euclidian distance between all pairs of points in the latent space of the given visit, clipped at the 90^th^ percentile, which prevented extreme outliers from unduely influencing cluster centriods. If subjects did not have at least four non-outlier trials per target after 7 blocks, additional calibration blocks were added automatically one at a time until the four-trial condition was met (no more than two such blocks were ever added). If subjects began their hand posture transition prior to being cued, did not initiate the transition within 0.5 seconds of the cue, or completed the transition more than 0.25 seconds prior to the training cursor coming to rest in the target, a corrisponding error message was displayed and that trial was repeated automatically (and data from the trial with the error was discarded).

After calibration, subjects freely explored an empty workspace with their cursor for 2 minutes before beginning a center-out target acquistion task (Supplementary Fig. 3C). A target appeared at the workspace center that subjects were required to reach and stay inside continuously for a 0.5 second hold period; we define a target “hit” as meeting this continuous reach and hold requirement, not merely the cursor reaching the target. The cursor was displayed as a dot approximately 25% of the area of the target, but only the cursor center was considered when determining target hits. Once the center target was hit, one of eight radially distributed peripheral targets was pseudorandomly displayed such that the order was not predictable and all targets appeared eight times (for a total of 8 targets x 8 occurences = 64 targets). Subjects then had 20 seconds to hit the peripheral target. After the 20 second timeout period or target hit, the centeral target reappeared and subjects were again required to reach and hold there to trigger the next trial, for which there was no timeout duration. After 150 minutes the task ended, whether or not subjects attempted all 64 peripheral targets, which happened 12 times in the 191 subject visits.

After the center-out task, subjects engaged in a typing task in which they were required to move and hold their cursor on keys displayed as a virtual keyboard to type target phrases (Supplementary Fig. 3D). To register as a keypress, the cursor needed to stay continuously within the key for 1s. A phrase would appear on the screen and subjects were instructed to type it as closely as possible within 2 minutes. The five phrases were (“Hello World”, “Cyber Glove”, “Potato Chips”, “Neuroscience”, and “Game Over”). Scoring was calculated as the Levenshtein distance (LD), i.e. how many character deletions, insertions, or substituions needed for the spelled phrase to match the given phrase of a trial. Values were then converted into a percent score, *S* = 100 · (*LD* − *c*)/*c*, where *c* is the total number of characters in the target phrase.

The workspace for the calibration, center-out, and keyboard tasks was a square with side length 2 a.u. (arbitrary units) centered at the origin, occupying 28.2 cm^2^ on the computer monitor. For the calibration (4 targets) and center-out task (8 targets), circular targets with a 0.15 a.u. radius were displayed at 0.85 a.u. from the the origin. The virtual keyboard had three rows of 10 square keys each, with a key side length of 0.198 a.u. that allowed a 0.01 a.u. gap between the edges of the keyboard and workspace boundery. The keys were laid out in a regular grid in standard QWERTY format, with the addition of “space” on row two and two “delete” keys and a “.” key on row three. We enforced that the cursor always remained inside the workspace bounderies by clipping all x and y positions. (Supplementary Fig. 3 – includes a diagram of experiment protocal and examples of all 3 tasks).

### Decoders

The five decoders used in this study generated cursor velocity commands every 50 ms. The velocity Kalman filter (vKF), the feedback intention-trained Kalman fitler (ReFIT), and direct regression with assisted calibration (DR-A) decoders were used only in the jaBCI model. The Wiener filter (WF) decoder was used only in the monkey-controlled iBCI. The descrete direction selection (DDS) decoder was used in both the jaBCI and iBCI experiments.

The vKF decoder updated cursor velocity at every time bin, *t*, according to

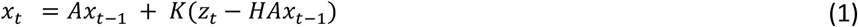

where *x* is the cursor state vector containing the positions, velocities, and offset, [*p_x_ p_y_ v_x_ v_y_* 1]^*T*^, *z* is the *c*-by-1 vector of neural firing rates (where *c* is the number of emulated neurons), *A* is the state transition model that predicts the next cursor state from the current state, *H* is the measurement model that generates the neural firing rates we expect given the state prediction, and*K* is the Kalman gain that weights the measurement estimates against the state estimates when creating the final update. The cursor position at *t* is set as the integrated velocity command produced by eq. 1, not the filter-predicted cursor position.

The state transition and measurement models were computed by regression on data from all non-outlier calibration trials, for *A* by relating the cursor state *x*_*t−*1_ to *x*_*t*_ and for *H* by relating *x*_*t*_ to *z*_*t*_. The Kalman gain was computed by sequentially stepping through each sample in the calibration data and making updates to error estimates and *K*. Terms *A, H, K*, the state transition noise matrix, the measurement model noise matrix, and the error estimates were computed as described in^15^. Terms *A, H*, and *K* were not updated during the closed-loop center-out task, effectively creating a constant gain Kalman filter.

The ReFIT decoder, described in detail elsewhere^16^, also applies cursor updates using eq. 1; however, it computes *A, H*, and *K* differently. Briefly, the regressions for *A* and *H* use inferred rather than actual cursor state data, *x*. That is, cursor velocity is assumed to be pointing toward the target and have zero speed when in the target regardless of the actual decoded state, and the state transition model assumes integrated cursor velocity predicts its subsequent position. Second, when computing *K* from calibration data the uncertainty propagated through each iteration is updated to reflect the fact that cursor position is known with complete accuracy (it is shown to the subject). This is achieved by setting to zero the position-related elements in the error estimate matrix in the standard Kalman update equations.

The DR-A decoder updated cursor velocity at every time bin, *t*, according to

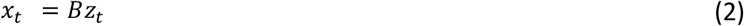

where *z* is the (*c* + 1)-by-1 vector of neural firing rates, *B* is the 2-by-(*c* + 1) matrix of regression coefficients mapping neural activity into cursor velocity (the additional row is for x and y velocity offsets), and *x*_*t*_ is the cursor state vector representing its velocit,y[*v_x_ v_y_*]^*T*^. The cursor position at *t* is set as the integrated velocity command produced by eq. 2. *B* is computed by regression relating *z*_*t*_ to *x*_*t*_ on data from all non-outlier calibration trials and using all emulated neurons with individual R^2^ with velocity at or above 0.03^17^. Additionally, during calibration the decoded cursor velocity was averaged with the training cursor velocity 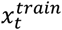 to produce the final decoded velocity *xt* at each time according to

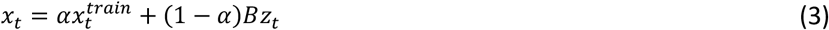

where (starting in block 3 when the decoded cursor first became visible alongside the training cursor) *α* decreased in each block from 0.8 by 0.2 to 0^17^. In cases where additional calibration blocks were added, *α* remained at 0 and all center-out and virtual keyboard trials had*α* = 0.

The WF decoder^49^ updated cursor velocity at every time bin, *t*, according to

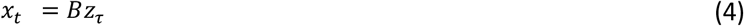

where *z*_*τ*_ = [*z*_*t*_ *z*_*t−*1_ … *z*_*t−τ*_ 1]^*T*^, *z* is a 1-by-*c* vector of neural firing rates, and the subscript denotes how many time bins in the past those firing rates occurred, with *t* being the current time. Thus, *z*τ is a concatenation of the firing rates of the *c* neurons across τ instants into the past and an offset term. As with DR-A, *x_t_* is the cursor state vector representing its velocity, [*v_x_ v_y_*]^*T*^, which gets integrated to determine the cursor position. Here *B* is the is the 2-by-(*cτ* + 1) matrix of coefficients mapping neural activity and its history into cursor velocity, which is computed by regression relating *z*_*τ*_ to *x*_*t*_ on data from all calibration trials. In the monkey iBCI task we used τ = 8, allowing WF to access firing rates up to 400 ms in the past. WF is like DR-A in that it is a single linear transformation mapping neural activity to cursor velocity with offsets to model baseline firing rates. It is dissimilar in that it includes 400ms of firing rate history when decoding velocity. Assisted calibration^17^ was not used in the monkey iBCI experiment.

The DDS decoder updated cursor velocity at every time bin, *t*, according to

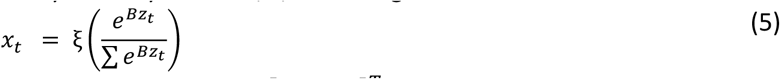

where *x*_*t*_, the current cursor state vector representing its velocity, [*v_x_ v_y_*]^*T*^ and *z_t_* is the (*c* + 1)-by-1 vector of neural firing rates (the additional row is for the offset term). *B* is the η-by-(*c* + 1) matrix containing the coefficients mapping the current neural activity into the multinomial logistic regression model probability estimates for each of the *η* selections. Meaning, at each time bin the multinomial model produces a set of *η* probabilities, 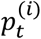, that represents its confidence that *z*_*t*_ coresponds to each of the *η* selections (the sum in eq. 5 runs over these *η* selections). Rather than choosing the velocity command with the multinomial model’s highest probability (winner take all), we performed a weighted average across the velocity commands of the most probable selections. The weighting was determined by a linear piecewise function of their probabilities, ξ(*p*_*t*_),

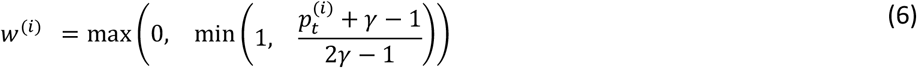

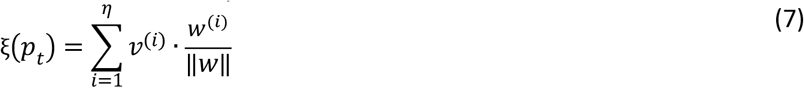

where 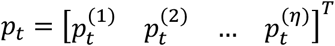 is the *η*-by-1 vector of multinomial model probabilities for each selection at time *t, v*^(*i*)^ is the velocity command of selection *i*, and *γ* ∈ (0.5 1] is a scalar “selection mixing” parameter. Equation 6 states that selections with probabilities below 1 − *γ* do not contribute to the average, a selection with probability greater than *γ* completely determines the result and forces all other weights to zero, and all selections with probabilities in between are weighted linearly in proportion to the magnitude of their probability. Equation 7 states that the vector of selection weights, *w*^(*i*)^, is renormalized to unity, then used in the weighted average of the velocities. *γ* = 1 produces an average weighted by the direct multinomial model probabilities, and as *γ* approaches 0.5 then ξ(*p*_*t*_) approaches a winner take all weighting. This formulation preserves the DDS philosophy of selecting from a short list of possible velocity commands, while also providing users with some visual feedback about when *z*_*t*_ nears a region in “neural activity space” that is close to a boundary between velocity command selections. We used *γ* = 0.85 and the *η* = 9 velocity selections [0 1], [0, 0.5], [1, 0], [0.5, 0], [0, −1], [0, −0.5], [−1, 0], [−0.5, 0], and [0, 0], where each corresponds to [*v_x_ v_y_*] in arbitrary workspace units per second.

DDS decoder calibration was done using the same task as all other decoders. To prepare the data for computing *B*, each trajectory from non-outlier trials was partitioned into fifths. Data from the beginning fifth of each trial was labeled as the “stop” selection, [0, 0]. Data from the middle fifth of each trial was labeled as the “fast” selection (1 unit/s) in the direction of the target (e.g., [0, 1] for the North target, [1, 0] for the East target, etc.). Data from the second and fourth fifth of each trial was labeled as the “slow” selection (0.5 units/s) in the direction of the target. After labeling, the multinomial logistic regression model, *B* (not including ξ(*p*_*t*_)), was fit using the scaled conjugate gradient descent algorithm.

### Analysis of Human Subject jaBCI Cursor Control

We used a full factorial ANOVA F Test design to test the effects of visit number and decoder on task performance, as well as any interactions between them. Post hoc tests were done using the Tukey HSD method. To control for learning to estimate the effect of neuron set on enter-out performance we fit a line to the number of target hits across visits, separately for each decoder group. We then subtracted this trend off each visit’s average target hits to remove the effect. All statistical analyses used a critical p-value of 0.05.

When analyzing the number of targets hit (Fig.1C) we considered only peripheral targets, not the return to the center target, which did not have a timeout period.

Time-to-target histograms were binned into 2-second intervals, normalized to unit area (Fig. 1D). Only successful trials were considered, creating a maximum at the 20 second timeout period. Group averages (Fig. 1E) were computed by averaging individual subject histograms in figure 1D.

To analyze cursor stopping behavior we computed normalized cursor speed histograms using all individual trials as observations, not averages (Fig. 2). Figure 2A shows histograms of cursor speed at the exact moment that a target hit was registered, thereby capturing speed at a single 50 ms bin. There were no cursor speed requirements to register a target hit, only that the cursor center remain completely within the target for the full 500 ms hold time.

The grid showing which DDS selections were mixed (Fig. 2C) is limited to showing only pairs of selections. Where more than two selections were mixed (≈7% of the time) the two selections with the largest weightings were counted. Data were pooled across all DDS subjects and visits (approximately 15 hours of recordings).

We computed the time directed toward target measure (Fig.2E) as the percentage of samples from a trial in which the direction of the decoded velocity was between the two rays emanating from the cursor position and contacting the target radius at the two opposing tangent points (see inset). All samples where the cursor was inside the target were counted as heading toward the target. All samples where the cursor was still in the home target from the previous trial were discarded, since often these involved heading corrections from the previous trial or reaction time. All successful trials were compiled into histograms of bin width 5% trial times and plotted separately for each decoder group.

Tortuosity, common in the vasculature literature^27,50^, was used as a measure of trajectory complexity and defined as the integrated instantaneous change in curvature, *κ*(*t*), normalized by total pathlength, *L*,

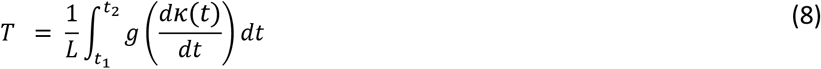

The curvature, *κ*(*t*), at each point is defined in the usual way as the reciprocal of the radius of the circle that best approximates the trajectory at each point. This measure is useful because it only penalizes direction changes if they happen with complex, continually changing arcs, meaning, there are ways to achieve low tortuosity besides straight-line trajectories. We take 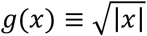 to enforce positive values and maintain numerical stability when computing *T* for trajectories with sharp corners (a case not encountered in blood vessels). We take *t*_1_ = 0.25s and *t*_2_ = 3s into the trial to consider the initial part of the trajectory that consists approximately of the subject’s first attempt. The first 0.25s often contained course corrections related to hitting the center home target at the end of the previous trial. We use at most 3 seconds of the trajectory to consider only the initial attempt at acquiring the target, and not the large corrections that would happen after the initial attempt failed (especially in the continuous velocity decoders), which would otherwise have inflated the tortuosity measures.

### Correlations Between Calibration and Cursor Control Outcomes in Human Subject jaBCI

We measured the difference between subjects’ decoded trajectories during calibration and the minimum jerk training trajectories shown during calibration to determine if it predicts closed-loop performance. Using each subject’s fully calibrated decoder from the center-out task, we decoded offline cursor kinematics from the emulated neural data they generated during the last 5 calibration blocks. Any automatically flagged calibration outliers were not used as part of the decoder calibration dataset, and were thus excluded here as well. We computed the discrete Fréchet distance between the decoded and training trajectories to produce a scalar dissimilarity measure. The discrete Fréchet distance is essentially the length of the longest line segment required to connect two points progressing along the paths through time. It is useful for trajectories because it does not harshly penalize similar shapes that are slightly out of phase. We then computed the correlation between the Fréchet distance for each subject-visit calibration and the number of targets they hit in the subsequent center-out task. Our inability to find a correlation between trajectory similarity and center-out performance was robust to many permutations of our analysis described above: i) the number of calibration trials used (all trials versus only later trials), ii) inclusion of outliers, iii) use of pointwise root mean squared error as a trajectory similarity measure, iv) correlating angular dispersion with center-out performance, v) correlating the average silhouette score of how trajectories through time clustered in a 2D PCA projection, and others.

We also tested whether highly separable neural population dynamics predicts closed-loop performance. To measure separability, we projected the neural trajectories from all calibration blocks into the top three PCs defined by all non-outlier calibration trials^51^. We then computed the pairwise discrete Fréchet distances between all neural trajectories in the reduced space. The distance matrix was used to compute the silhouette score for each trajectory belonging to the cluster corresponding to the calibration target. The silhouette score is the normalized difference between the average distance of a trajectory to those in its nearest neighboring cluster and the average distance of that trajectory to all others in its own cluster. Thus, larger values indicate greater separation with (distance from) trajectories of neighboring clusters than its own cluster, making the data more separable. We then computed the correlation between the average silhouette score^48^ for each subject-visit calibration and the number of targets they hit in the subsequent center-out task.

### Protocol for Monkey iBCI Use

We used one 14-kg adult male rhesus monkey (Macaca mulatta) in this study, who had been previously trained, and implanted with a 128-channel “Utah” electrode array (Blackrock Neurotech, Inc.) in the arm/hand representation of the left primary motor cortex (M1). The monkey performed an isometric wrist task requiring him to move a cursor from the center of the monitor to one of eight radial targets by exerting forces on a small box placed around his right hand (Fig. 3A). The box was padded to comfortably constrain the monkey’s hand and minimize its movement, and the forces were measured by a 6-DOF load cell (JR3 Inc, CA) aligned to the wrist joint. Flexion/extension force moved the cursor right and left respectively, while force along the radial/ulnar deviation axis moved it up and down. Each trial started with the appearance of a center target which the monkey was required to hold for a random time (0.2 - 0.5 s), after which one of the outer targets, selected in a block-randomized fashion appeared, accompanied by an auditory go cue. The monkey needed to move the cursor to the target within 2.0 s and hold the cursor within the target for 0.3 s to receive a liquid reward.

For each experimental session, M1 activity, and force were recorded using a Cerebus system (Blackrock Neurotech, Inc). The neural signals on each channel were digitalized, bandpass filtered (250 - 5000 Hz) and converted to spike times based on multi-unit threshold crossings (5.5 times the root-mean-square amplitude of the signal on each channel). We computed the spike counts in 50 ms, non-overlapping bins to obtain an estimate of firing rate as function of time for each channel.

At the start of each of the seven recording sessions, we created a calibration dataset as the monkey performed the isometric wrist task in “hand-control” mode (Fig. 3A) for 10 minutes. We used multiunit activity recorded from M1, along with cursor velocity to compute both Wiener filter (WF) and DDS decoders, using data from successful trials limited to the interval between the go cue and reward. To calibrate the DDS decoder, cursor velocities were divided into the same 9 selections described in the jaBCI study.

All surgical and experimental procedures were approved by the Institutional Animal Care and Use Committee (IACUC) of Northwestern University under protocol #IS00000367 and are consistent with the Guide for the Care and Use of Laboratory Animals.

## Acknowledgements

We acknowledge Maral Daneshyan, Paulwin Arancherry, and Hary Usaquen for assistance with data collection, and Steafan Khan for discussions about decoder design. This work was funded by the NIH R01NS109257.

## Author Contributions

Danziger created the experiment design, Alcolea ran experiments with human subjects and analyzed the data. Ma ran experiments with the monkey, collected the data, analyzed the results, and added the results and methods of monkey analysis to paper. Bodkin designed the infrastructure used to run the monkey experiments and analyzed the results. Miller supervised and guided the steps required to complete the experiment. Writing was primarily done by Alcolea and Danziger, and Miller had major input into the writing process and structure of paper.

## Supplemental Figures

**Supplemental Figure 1:**
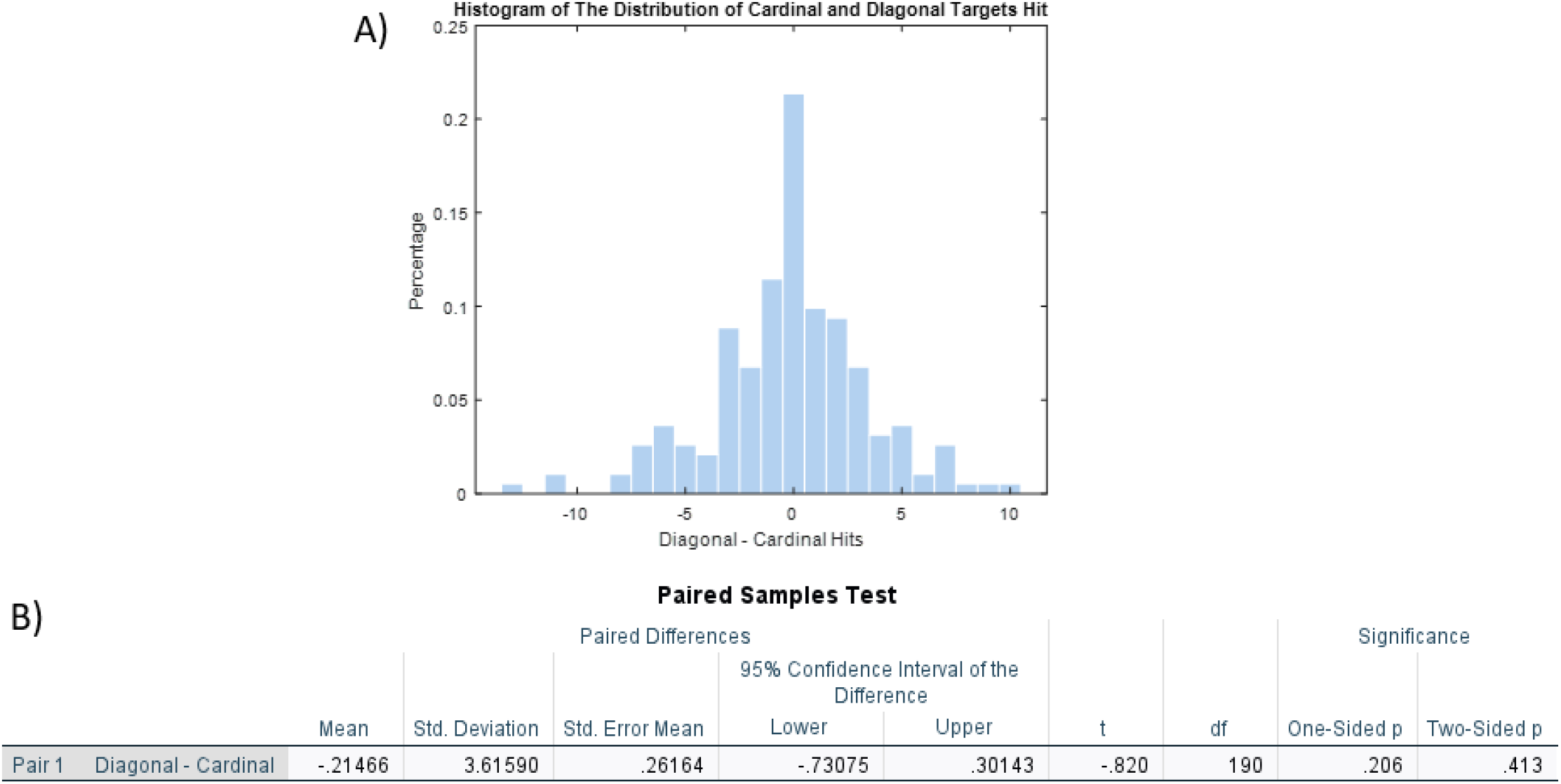
Cardinal vs. diagonal target hit analysis. A) This histogram shows the distribution of the difference between diagonal targets and cardinal targets hit in the CenterOut task for all subject-visits. B) Analysis of histogram and additional statistical analysis with a two-sided p score of 0.413 showed that subjects had no preference in hitting cardinal over diagonal targets across all subject visits. This supports the use of only 4 directions during calibration.

**Supplemental Figure 2:**
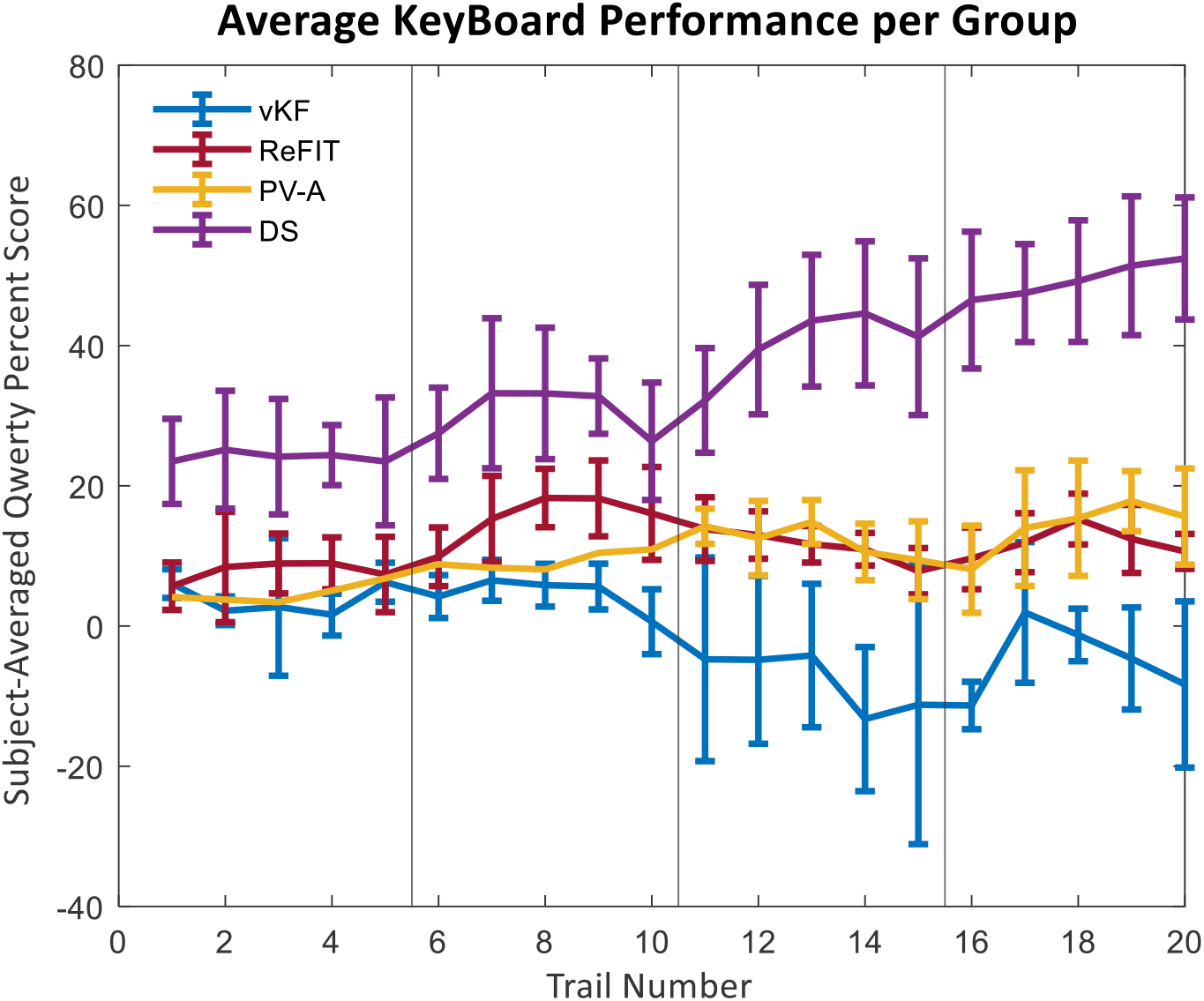
Performance on the virtual QWERTY keyboard task. Each line is the mean percent score (y-axis) of every trail. Vertical lines denote transitions between visits. Error bars in each line represent the standard error of all subjects in a group. This task was considerably more difficult than the center-out task, with only DDS subjects able to meaningfully type (note scores of near zero for ReFIT and DR-A, and a negative score for vKF). Negative scores occur when the subject types more incorrect keys than correct ones, meaning vKF users would have performed had they not typed at all.

**Supplemental Figure 3:**
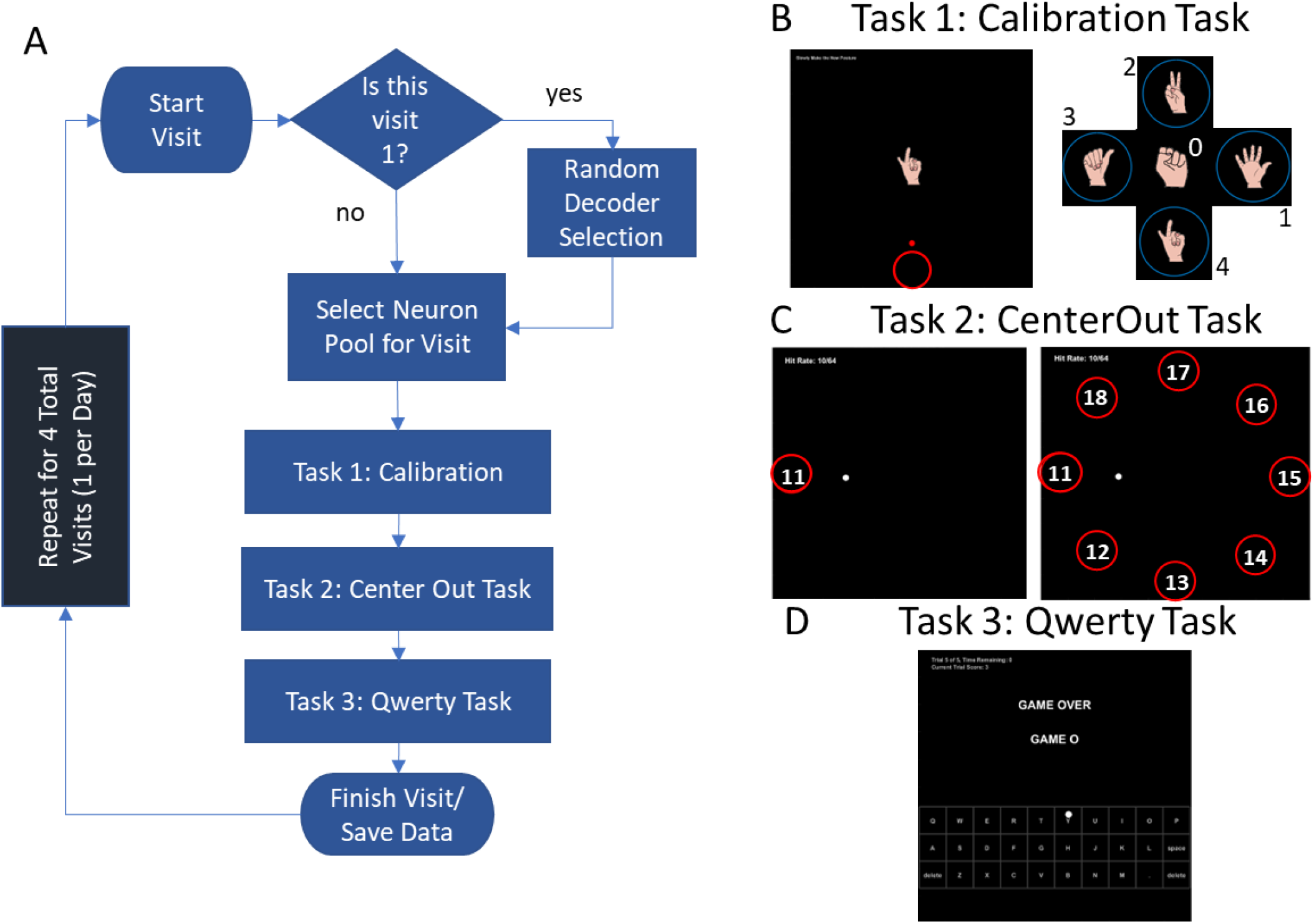
Experimental protocol. A) Describes the protocol for a subject’s 4 visits. In visit 1 subjects will be randomly placed into a decoder group, will be selected into the visit 1 emulated neuron set, and complete 3 tasks. In the following visits (held on different days) subjects will be placed into different emulated neuron sets and complete the same 3 tasks. B) Depiction of the calibration task, showing the four displayed postures and their corresponding target direction. C) Depicts the Center-Out task, by showing how a singular trail would look like if a subject got the left target in trial 11 (left-B), and showing all target locations that users will have to target throughout the 64 trials (right-B). Each target will be appearing 8 times throughout the task. D) Depiction of Qwerty Task, which involves subjects spelling out 5 words or phrases in 2 minutes or less each.

**Supplemental Table 1.** Statistical analysis of between group effects based on the results of the CenterOut task using the Full Factorial Anova F Test. Shows the effect of visits and the interaction between days and decoders. Results suggest that decoders and visits have a significant effect with a p < 0.001, while exhibiting no significant interaction. Post hoc analysis with Tukey HSD further revealed that all decoders are significantly different from one another and that only visits 1 and 4 are significantly different.

| Source | Type III Sum of Squares | df | Mean Square | F | Sig. |
| --- | --- | --- | --- | --- | --- |
| Corrected Model | 52081.770<sup>a</sup> | 15 | 3472.118 | 27.705 | <.001 |
| Intercept | 225220.493 | 1 | 225220.493 | 1797.104 | <.001 |
| Groups | 49119.354 | 3 | 16373.118 | 130.646 | <.001 |
| Visits | 2429.418 | 3 | 809.806 | 6.462 | <.001 |
| Groups * Visits | 521.774 | 9 | 57.975 | .463 | .898 |
| Error | 21931.727 | 175 | 125.324 |  |  |
| Total | 299389.000 | 191 |  |  |  |
| Corrected Total | 74013.497 | 190 |  |  |  |

**Supplemental Table 2.** Statistical analysis of between group effects based on the results of the CenterOut task using the Full Factorial Anova F Test shows the effects of neuron set and the interaction between neuron set and decoder. The reason the tested effects change is due to the fact learning has been removed to focus on the effect of the different neurons used in each visit. Analysis revealed that there was no effect of neuron set on subject performance and that even when learning is removed the effect of decoders remained significant. Important value to consider is the neuron set p of 0.722, which is greater than 0.05 suggesting no significant difference of performance based on the neuron set.

| Source | Type III Sum of Squares | df | Mean Square | F | Sig. |
| --- | --- | --- | --- | --- | --- |
| Corrected Model | 51694.780<sup>a</sup> | 15 | 3446.319 | 27.070 | <.001 |
| Intercept | 166831.281 | 1 | 166831.281 | 1310.417 | <.001 |
| Groups1 | 51258.688 | 3 | 17086.229 | 134.208 | <.001 |
| Neurons | 240.150 | 3 | 80.050 | .629 | .597 |
| Groups1 * Neurons | 185.046 | 9 | 20.561 | .161 | .997 |
| Error | 21897.591 | 172 | 127.312 |  |  |
| Total | 240973.287 | 188 |  |  |  |
| Corrected Total | 73592.371 | 187 |  |  |  |

**Supplemental Table 3.** The table shows a One-Way Anova F-Test analysis testing the effect of decoder groups after removing all learning gained from subjects throughout the visits. This analysis helps show how the sum of squares between and within groups differ. Analysis shows that despite the large variance of subjects within groups, the difference between groups was larger.

|  | Sum of Squares | df | Mean Square | F | Sig. |
| --- | --- | --- | --- | --- | --- |
| Between Groups | 51377.316 | 3 | 17125.772 | 140.546 | <.001 |
| Within Groups | 22908.125 | 188 | 121.852 |  |  |
| Total | 74285.441 | 191 |  |  |  |

